# Causal Modeling Reveals Cell-Cell Communication Dynamics in the Tumor Microenvironment During Anti-PD-1 Therapy in Breast Cancer Patients

**DOI:** 10.1101/2025.10.08.681192

**Authors:** Aodong Qiu, Han Zhang, Joseph D Ramsey, Bryan Andrews, Boyang Sun, Shuangxia Ren, Mengyao Lu, Kun Zhang, Gregory Cooper, Binfeng Lu, Lujia Chen, Xinghua Lu

## Abstract

**Background:** Immune checkpoint blockade (ICB) targeting PD-1/PD-L1 plays a crucial role in breast cancer treatment. Despite clinical success, how ICB reshapes the tumor microenvironment (TME) to enhance anti-tumor activity remains unclear. Anti-PD-1 therapy alters TME cells beyond PD-1+ T cells, with extensive cell-cell communication (CCC) playing a key role. Understanding the dynamic CCC changes upon anti-PD-1 treatment can illuminate ICB mechanisms of action and TME dynamics.

**Methods:** We analyzed single-cell RNA-seq data from 31 breast cancer patients before and after anti-PD-1 (pembrolizumab) treatment (Bassez et al., 2021). We identified differentially expressed genes (DEGs) induced by treatment in major cell types. We then applied an instrumental variable approach to uncover causal relationships between T-cell and non-T-cell DEGs. We further mapped ligand-receptor interactions mediating signal transduction between cells and constructed a CCC network from T to non-T cells.

**Results:** Anti-PD-1 therapy induced widespread transcriptional changes across multiple cell populations. Key pathways modulated in T cells included NF-κB, interferon-γ, and interleukin signaling. CD4+ and CD8+ exhausted T cells engaged in distinct ligand-receptor interactions with tumor-associated macrophages (TAMs) and other types of cells, reshaping the TME. Our results indicated CD4+ exhausted T cells activated M1-like TAMs via TNF-TNFRSF1A and TNFSF14-LTBR, while CD8+ exhausted T cells engaged M1-like TAMs through ICAM1-ITGAL/ITGB2 and CCL8-CCR2, promoting anti-tumor immunity. Conversely, immunosuppressive interactions were also observed, such as TNF–TNFRSF1B (TNFR2) and TNFSF14–TNFRSF14 (HVEM) from CD4⁺ T cells, as well as CSF1–CSF1R and RPS19–C5AR1 from CD8⁺ T cells, which likely promote M2-like tumor-associated macrophage (TAM) polarization and contribute to pro-tumor immune regulation and resistance to therapy. Notably, key receptors in the causal CCC networks, such as C5AR1, TNFR2, and CSF1R, emerged as potential targets to enhance anti-PD-1 efficacy.

**Conclusions:** These findings elucidate TME remodeling during anti-PD-1 therapy and underscore the pivotal role of CCC in treatment response. Our study identifies critical communication networks that may be biomarkers for immunotherapy responsiveness and highlights novel therapeutic targets, including C5AR1 and HVEM. Furthermore, our application of causal inference methodologies provides a robust framework for dissecting CCC mechanisms in immunotherapy.

## Introduction

Breast cancer remains the most common malignancy affecting women worldwide, characterized by diverse biological behaviors and responses to treatment across its subtypes. This variability presents significant challenges in developing universal therapeutic strategies^1^. Among breast cancer subtypes, triple-negative breast cancer (TNBC) is notably aggressive and lacks targeted therapies due to its absence of estrogen and progesterone receptors and HER2 expression^2^. Historically, breast cancer has not been considered highly immunogenic, but in recent decades, discoveries have shown substantial tumor-infiltrating lymphocytes (TILs) in some subtypes, especially in TNBC, suggesting potential responsiveness to immunotherapies^3–5^.

Immune checkpoint blockade (ICB) therapies have revolutionized cancer treatment by targeting regulatory pathways in T cells to enhance the immune system’s response against cancer cells^6^. Agents targeting the Programmed Death-1 (PD-1) pathway, which is often exploited by tumors to evade anti-tumor immune responses, have shown promising results in various cancers, including breast cancer^7^. The immunotherapy response rate in breast cancer remains modest. In TNBC, the most immunogenic subtype, the response rate to anti-PD1 monotherapy is generally below 20%^8^. Despite their clinical success, the precise mechanisms by which ICB therapies modify the TME and promote anti-tumor activity remain incompletely understood. The complexity of the TME, which includes cancer cells, immune cells, stromal cells, the extracellular matrix, and soluble factors, makes it difficult to fully comprehend how these therapies alter cellular communication and what the mechanisms of treatment resistance are^9^.

Recent advances in computational methods have enabled the systematic inference of CCC networks from single-cell transcriptomic data. Tools such as CellChat^10^, NicheNet^11^, and CellPhoneDB^12^ have been widely applied to characterize ligand-receptor-mediated signaling across diverse cell types. However, most existing approaches are based on correlation analysis and do not infer causal relationships between interacting cells. This highlights the need for causal inference to uncover communication channels and their impact on cellular states that may underlie therapeutic response or resistance.

This study investigated the CCCs within the TME of breast cancer patients undergoing anti-PD-1 therapy. By analyzing single-cell RNA sequencing data from pre- and on-treatment conditions from 31 patients receiving anti-PD-1 treatment reported by Bassez, *et al*^13^, we examined the changes in the cellular state of diverse cells. Further, we investigated the mechanisms underlying the coordinated changes among cells within TME. Employing principled causal inference methodology, we constructed a comprehensive map of the cell-cell communication network influenced by anti-PD-1 therapy. We further identified pivotal ligand-receptor pairs that likely mediate communication among various cell types.

Our analyses provide a detailed understanding of the dynamic interplay of cells within the TMEs during anti-PD-1 therapy, providing new information on the mechanisms of action of anti-PD-1 therapy. This deeper knowledge could facilitate the identification of new biomarkers for predicting heterogeneous treatment responses to anti-PD-1 regimens, potentially enhancing the design and customization of immunotherapeutic strategies for breast cancer patients. Finally, this study demonstrated the utility of causal inference methodology for mechanistic studies of CCC.

## Results

### Profile of cell landscape under pre- and on-treatment conditions

To evaluate the impact of anti-PD-1 therapy (pembrolizumab), we analyzed single-cell and TCR-seq data from a window-of-opportunity study reported by Bassez, *et al*^13^. The dataset included 168,970 cells from 31 patients, collected under pre- and on-treatment conditions. The data were processed following a quality control protocol, including PCA, neighbor detection, and Uniform Manifold Approximation and Projection (UMAP) algorithms. Differentially expressed genes (DEGs) in response to treatment were identified in different cell types (or cell clusters), laying a foundation for studying CCC-induced coordinated changes among different cells by the treatment. We then employed causal inference methods, including the instrumental variable model and conditional independence tests, to build a CCC network. Figure 1a illustrates the workflow of this study.

**Figure 1.**
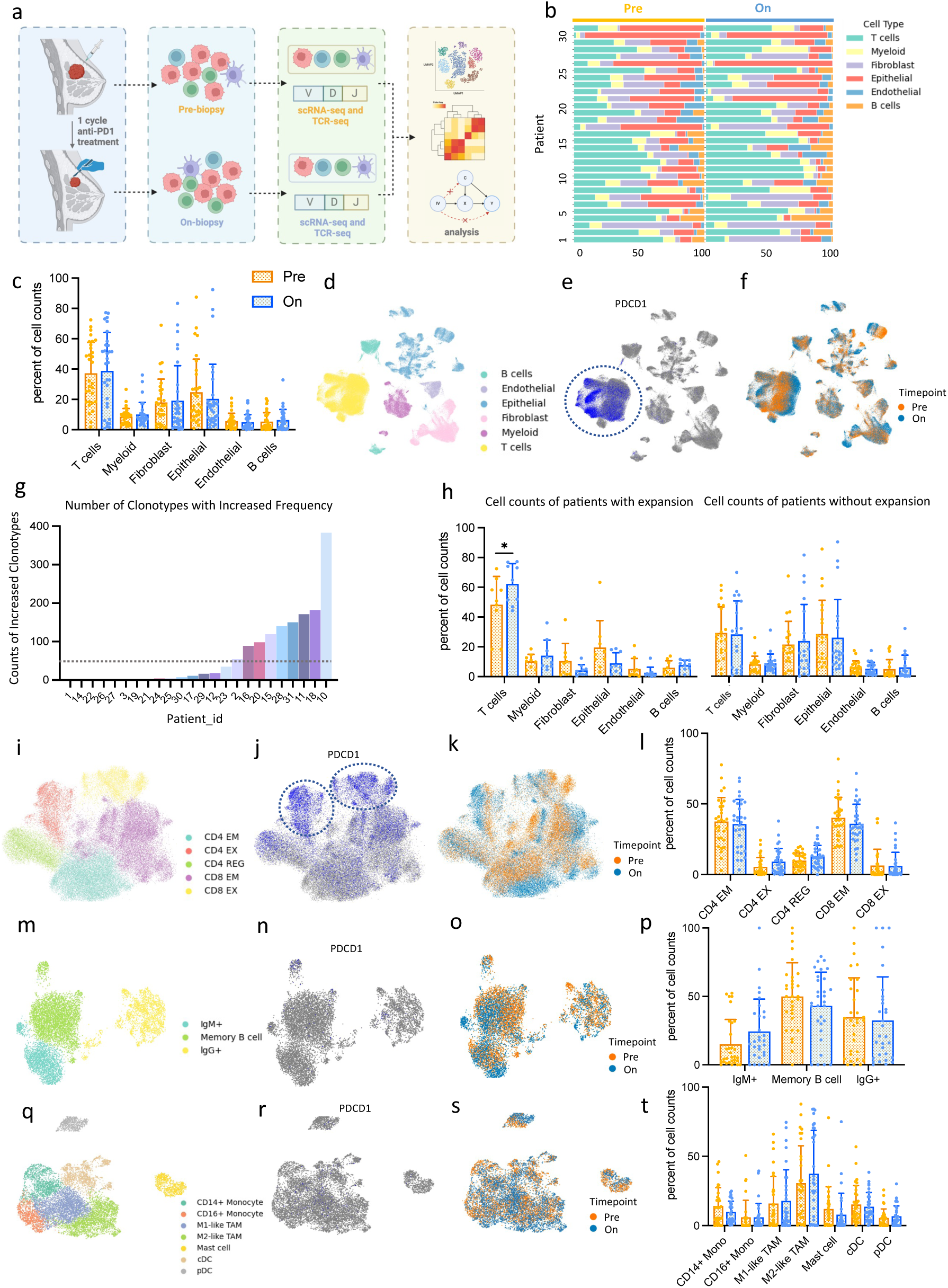
Overview of experiment process and profile of pre- and on-treatment landscape. **a**, Overview of the data collection and experiment design. Breast cancer biopsies are collected before and after one cycle of anti-PD-1 treatment (pembrolizumab) from 31 patients. Single-cell RNA sequencing data were collected from 168,970 cells. Then, a series of data analyses were performed to build a causal CCC network. **b**, Proportions of each cell type in each sample. **c**, Percentage of cell counts for each cell type in the samples before or after treatment. **d**, UMAP of 168,970 cells colored by cell types. **e**, UMAP colored by the expression level of *PDCD1*. **f**, UMAP colored by the time point of treatment. **g**, Number of clonotypes with increased frequency given TCR-seq data. **h**, Percentage of cell counts for each cell type in the samples with or without clonal expansion. **i-l**, UMAP of T cells colored by cell subtypes, expression level of *PDCD1, and* time point. Percentage of cell counts for T cell subtypes in the samples before or after treatment. **m-p**, UMAP maps of B cells colored by cell subtypes, timepoint, and expression level of PD1. Percentage of cell counts for B cell subtypes in the samples before or after treatment. **q-t**, UMAP maps of myeloid cells colored by cell subtypes, timepoint, and expression level of PD1. Percentage of cell counts for myeloid cell subtypes in the samples before or after treatment.

We inspected the landscape of cells before and after treatment. The cellular composition is highly diverse among the tumor TMEs from 31 patients, potentially due to inter-tumor heterogeneity in TEMs or sampling different areas of tumors in different patients (Figure 1b). However, the overall cell counts across various cell types remained largely unchanged throughout the treatment course (Figure 1c). Cells of distinct lineage formed clusters along the cell types, including B, endothelial, epithelial, fibroblasts, myeloid, and T cells (Figure 1d). For each cell type, a notable shift was observed on the UMAP map after treatment (Figure 1f). The position shift of a group of cells on the UMAP map indicates that the treatment has significantly altered the gene expression profiles of these cells. Theoretically, the anti-PD-1 treatment only targets the cells expressing PD-1^14^. Since *PDCD1* (encoding PD-1) expression was predominantly observed in T cells, with negligible expression in other cell types (Figure 1e), we hypothesized that the transcriptomic changes induced by the treatment resulted from CCC between the T cells expressing PD-1 and other cells.

Based on T cell receptor sequencing (TCR-seq) data, T-cell clonal expansion was observed in 9 patients; 20 patients showed no T-cell expansion; 2 patients lacked TCR-seq data, leaving their clonal status undetermined (Figure 1g). In patients with expansions (Es), the composition of cell types changed more obviously after treatment compared to the patients with no expansions (NEs), and T cells significantly increased after treatment (paired t-test, p < 0.05) (Figure 1h).

T cells were clustered into five major subtypes, including CD4+ effector/memory cells (CD4 EM), CD4+ exhausted cells (CD4 EX), CD4+ regulatory cells (CD4 REG), CD8+ effector/memory cells (CD8 EM), and CD8+ exhausted cells (CD8 EX) (Figure 1i). Despite the T cell UMAP plot having an obvious shift after treatment, the counts of T cell subtypes remained stable (Figure 1l). *PDCD1* expression was broadly distributed among T cells, with a high concentration in CD4 EX and CD8 EX (Figure 1j). B cells are divided into IgM+, Memory, and IgG+ cells (Figure 1m), whereas myeloid cells are classified into CD14+ monocytes, CD16+ monocytes, M1-like tumor-associated macrophages (TAMs), M2-like TAMs, mast cells, plasmacytoid dendritic cells (pDC) and conventional dendritic cells (cDC) (Figure 1q). Both B and myeloid cells have almost no *PDCD1* expression, and the subtype composition remained consistent after anti-PD1 treatment. (Figure 1n, p, r, t)

### Anti-PD-1 treatment induced broad gene expression changes in diverse cell populations

We pooled cells belonging to a common lineage, e.g., T cells, from pre- and on-treatment samples for each patient. We constructed pseudo-bulk RNA expression profiles under these conditions, which enabled us to detect gene expression changes in response to the treatment using paired t-tests. T cells expressing *PDCD1* are likely the target cells of the treatment. We pooled cells expressing *PDCD1* and performed DEG analysis (Figure 2a). The top 5 induced DEGs based on the p-value include *PRDM1*, *TXNIP*, *TSC22D3*, *FKBP5*, and *NFKBIA*, whereas the top 5 suppressed DEGs are *ZFP36L1*, *IRF1*, *FASLG*, *TNFSF14,* and *IER2*. *PRDM1* is an important transcription factor involved in T and B cell differentiation. We noted that the induction of *PRDM1* by anti-PD-1 treatment in breast cancer is associated with the changed expression status of its known downstream target genes, including *IL10*, *IL2*, and *BATF*^15^. Another interesting finding is that many genes regulated by NF-κB are significantly differentially expressed in response to anti-PD-1 treatment, including *NFKBIA*, *CXCR4*, *FASLG*, *TNFAIP3*, etc.^16^, which indicates a change in the state of this pathway upon treatment.

**Figure 2.**
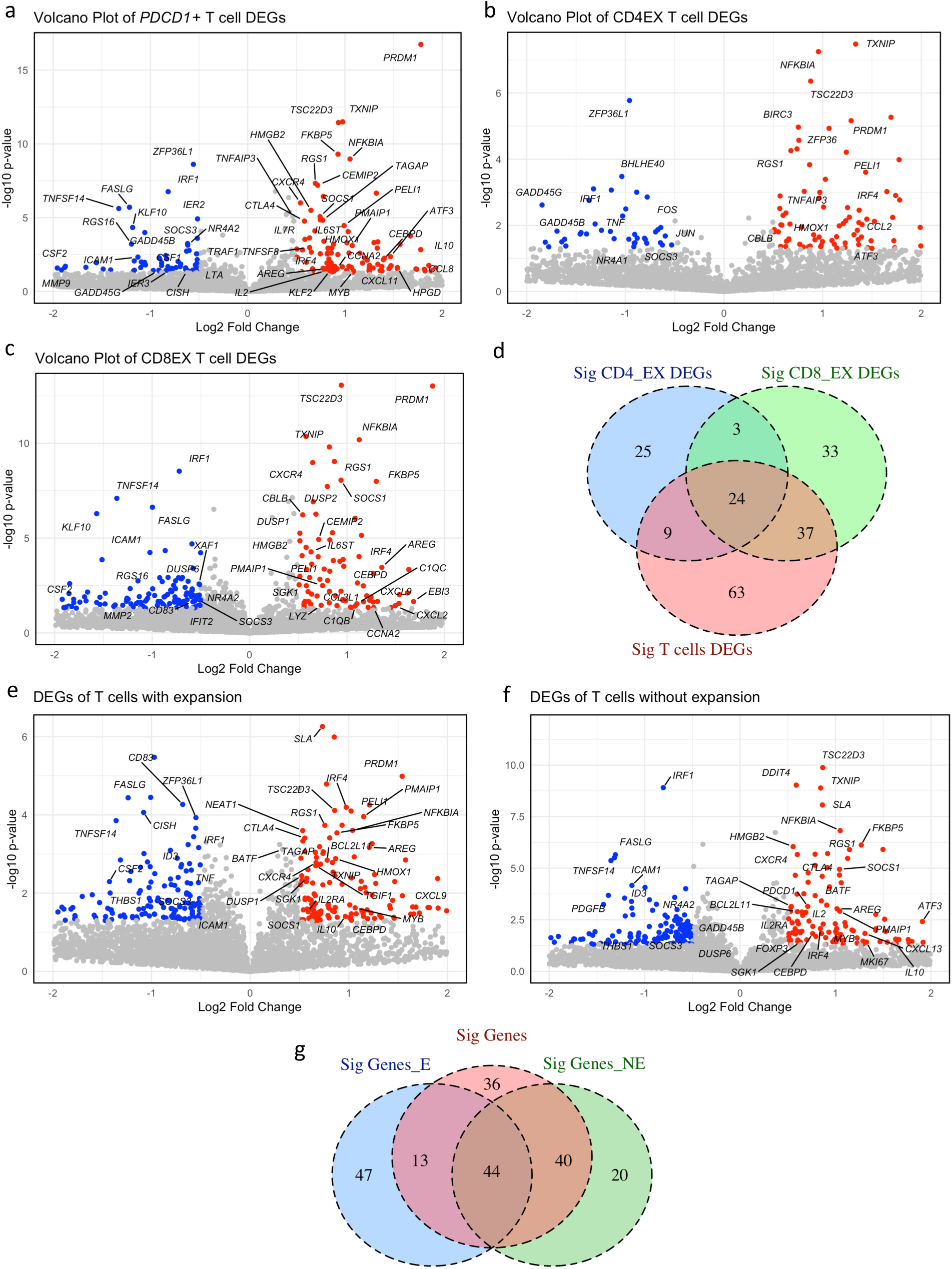
Significant DEGs after the anti-PD-1 treatment in T cell subclusters. **a**, Volcano plot showing DEGs in T cells expressing *PDCD1*, comparing before treatment (N = 5,610 cells) and after treatment (N = 8,457 cells). Dots on the volcano plot: Gray, P value is not significant; Red, P < 0.05 and log2 fold change is more than 0.5; Blue, P < 0.05 and log2 fold change is less than -0.5. P values were obtained by the paired t-test. **b**, Volcano plot showing DEGs in CD4+ exhausted T cells comparing before treatment (N = 1,932 cells) and after treatment (N = 3,746 cells). **c**, Volcano plot showing DEGs in CD8+ exhausted T cells comparing before treatment (N = 3,000 cells) and after treatment (N = 2,365 cells). **d**, Venn diagram comparing significantly DEGs across CD4+ T cells, CD8+ T cells, and all PD1+ T cells. Overlapping regions indicate genes shared between these groups, while non-overlapping regions represent group-specific DEGs. **e,** Volcano plot showing DEGs in T cells with expansion comparing before treatment (N = 11,724 cells) and after treatment (N = 18,414 cells). **f,** Volcano plot showing DEGs in T cells without expansion, comparing before treatment (N = 10,405 cells) and after treatment (N = 9,512 cells). **g**, Venn diagram comparing significant DEGs of T cells in patients with T cell expansion, without T cell expansion, and all T cells.

Investigation of distinct responses in different subpopulations of T cells can reveal the detailed roles of these cells in modulating TME. We examined and compared the DEGs from exhausted CD4+ and CD8+ T cells (Figure 2b, c), which expressed high levels of *PDCD1* compared to other T cell subtypes. The DEGs among these two important T cell subpopulations did not completely overlap (Figure 2d). While they share common DEGs, such as *PRDM1*, *NFKBIA*, *TXNIP*, and *TSC22D3*, they also have specific DEGs that are not differentially expressed in the other types of cells, such as *SESN1*, *KLF2*, and *BHLHE40* in CD4+ exhausted T cells and *CEBPD*, *TNFSF8*, and *SOCS1* for CD8+ exhausted T cells. Interestingly, *CTLA4* was induced in CD4+ exhausted T cells by anti-PD-1 treatment, suggesting potential interactions between PD-1 and CTLA-4 signaling, as reported in previous studies ^17^.

Based on T cell clonal expansion following anti-PD-1 treatment, tumors were classified as E (with clonal expansion) and NE (no clonal expansion), reflecting differential T cell responses across distinct TMEs. Although T cells from both tumor subgroups shared a set of DEGs in response to treatment, E and NE tumors also exhibited distinct DEG signatures, suggesting divergent transcriptional responses to the same immunotherapy depending on the TME context (Figure 2e–g). Some upregulated DEGs from E tumors involve anti-tumor immune responses, such as *DUSP1*^18^ and *HMOX1*^19^. Interestingly, *CTLA4* is also among the DEG in E tumors, again suggesting an interaction of PD-1 and *CTLA4* signaling pathways.

### Discovering potential cell-cell communication between PD1+ T cells and non-T cells in TME through causal analysis

Our analyses revealed a significant number of DEGs among non-T cells in response to anti-PD-1 treatment. Furthermore, many non-T-cell DEGs exhibited high correlations with some T-cell DEGs, which indicates possible CCC. We denote a DEG from PD1+ T cells as *X* and a DEG from non-T cells as *Y*. We performed a series of correlation analyses among the top-ranking DEGs from PD1+ T cells and those from non-T cells to search for significantly correlated *X-Y* pairs. The correlations between the top 20 DEGs from T and myeloid cells are shown as a heatmap in Figure 3a. Interestingly, certain *X-Y* pairs share a high Pearson correlation (|correlation| > 0.7), indicating that anti-PD-1 therapy induced coordinated changes in multiple cell types in TME.

**Figure 3.**
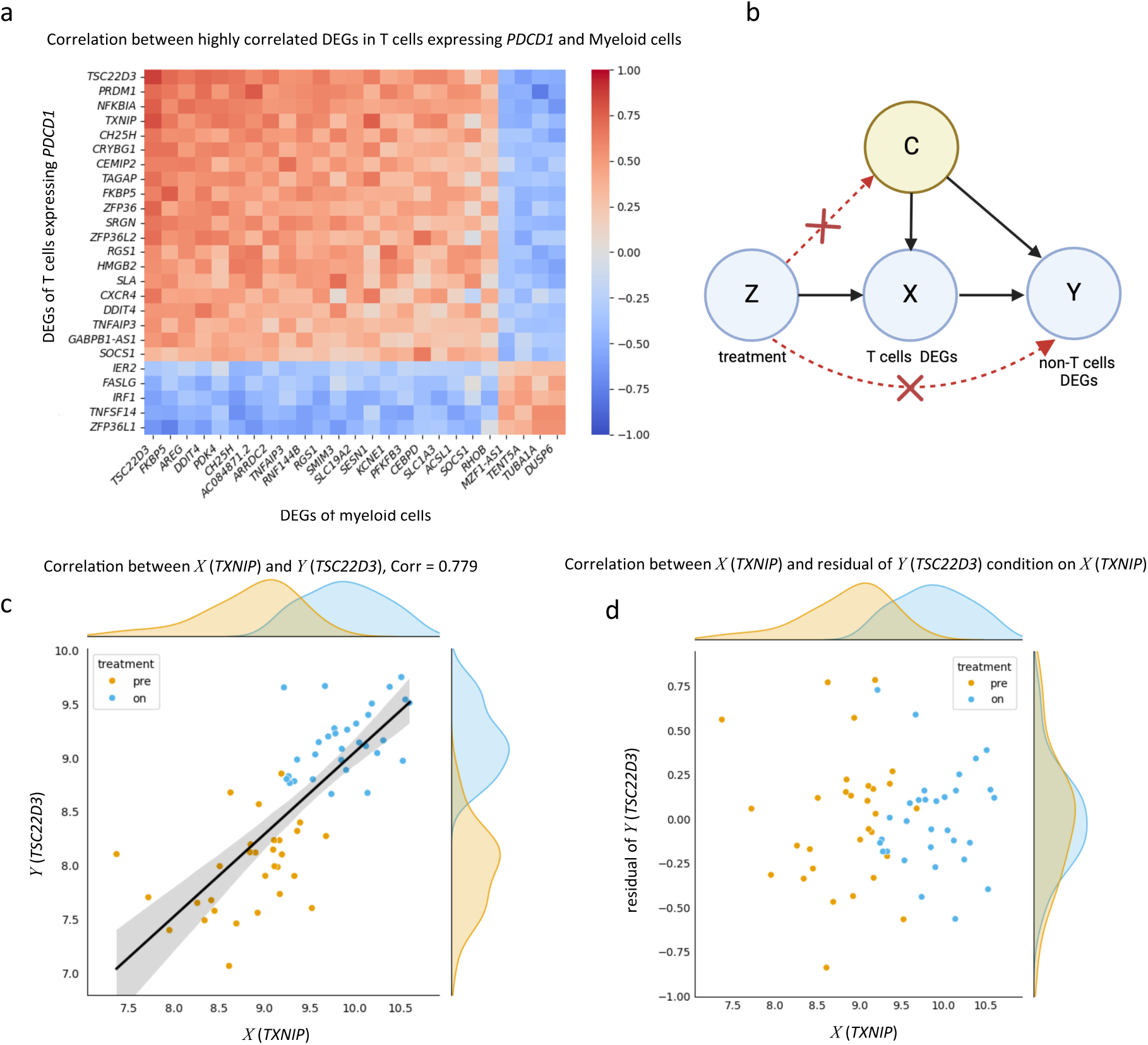
Significant DEG correlation between specific T cell clusters and other cell type clusters. **a**, Heatmap of correlation between highly correlated DEGs between T cells expressing *PDCD1* and Myeloid cells. **b**, Causal graph of application of the instrumental variable. **c**, The scatter plot showing the correlation between *X* (*TXNIP*) and *Y* (*TSC22D3*). Pearson correlation = 0.779. **d**, The scatter plot showing the correlation between *X* (*TXNIP*) and the residual of *Y* (*TSC22D3*) condition on *X* (*TXNIP*).

Since myeloid cells do not express *PDCD1* (PD-1), the anti-PD-1 treatment is unlikely to affect myeloid cells directly. Thus, it can be hypothesized that such a strong correlation between DEGs from T and myeloid cells is likely induced by CCCs. In other words, changes in the T cell state (reflected by a surrogate variable *X*) can causally influence the state of myeloid (reflected by a surrogate variable *Y*) through CCC. To test this causal hypothesis, one must first exclude the scenario that the correlation between *X* and *Y* is due to confounding by unobserved confounders. Here, we employed a principled causal analysis method called the instrumental variable (IV) estimation^20^ (Figure 3b) to test whether the correlation between *X* and *Y* is due to CCC in response to anti-PD1 treatment.

Here we use the example of anti-PD-1 treatment (*Z*), *TXNIP* (a DEG from T, *X*), and *TSC22D3* (a DEG from myeloid, *Y*) to illustrate the IV test framework. Our prior knowledge of the lack of PD-1 expression in myeloid excludes the possible causal edge from *Z* to *Y*; as an exogenetic perturbation, *Z* is unlikely to be connected with the latent confound *C*. We applied the IV test to examine whether the correlation between *X* and *Y* is mediated via CCC between cells. *TXNIP* expression is correlated with *TSC22D3* (Pearson correlation = 0.779), and both have a bimodal distribution associated with treatment condition (Figure 3c). After conditioning on *TXNIP* expression, *TSC22D3* expression is independent of treatment (Figure 3d). The result supports the hypothesis that the treatment influences the gene in myeloid through CCC between PD-1+ T cells and myeloid. More details regarding the molecular mechanism underlying this CCC channel will be discussed in a later section.

We systematically performed the IV analysis to investigate potential causal relationships between the signal regulating DEGs of PD1+ T cells and non-T cells subtypes (Supplementary Table S1). Comparing the DEG pairs passing the IV test (p-value < 0.05, q-value < 0.1), we noted that PD1+ T cell DEGs tend to exhibit higher correlations with myeloid and endothelial DEGs but relatively lower correlations with fibroblast and B cells (Supplemental Figure 1a, b and Supplemental Figure 2a, b). We also did the same analysis on the subtypes of PD1+ T cells and non-T cells, such as the CD4+ exhausted T cells and Monocytes (Supplemental Figure 3a, b).

### Building CCC networks by searching for potential ligand-receptor interactions

We hypothesize that anti-PD-1 treatment modulates TME by directly affecting PD1+ T cells first and then non-T cells through CCC (Figure 4a). The IV analyses reveal such potential CCC between PD1+ T cells and other cell types by excluding possible confounding factors (Figure 4b). In recent studies, the concentration of a ligand-receptor complex (LR) in a tumor sample has been computed using RNA expression levels of the ligand and receptor as surrogates for their protein concentrations. We adopted a similar approach (see Methods) and represented an LR complex in a causal graph (Figure 4b). From a causal analysis viewpoint, if an LR pair mediates the signal transduction between two cells and leads to correlated *X* and *Y*, conditioning on its state would render the variables *X* and *Y* independent, i.e., conditional independence (Figure 4b).

**Figure 4.**
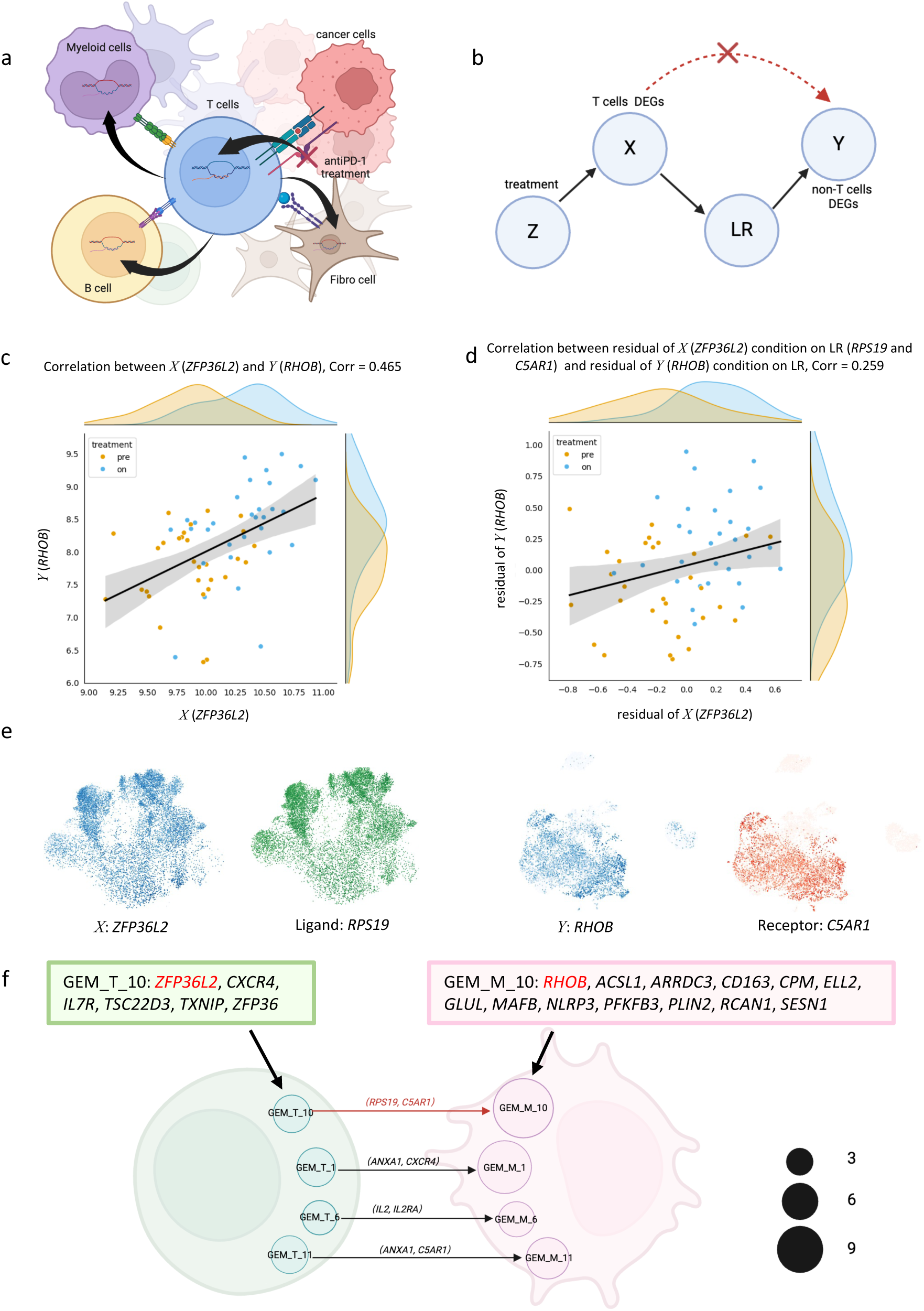
Validate the significant DEG correlations between T and non-T cells using causal inference methods. **a**, A simplified illustration of the CCC network after anti-PD1 treatment. **b**, The causal graph of application of the conditional independence test. **c**, The scatter plot depicting the correlation between *X* (*ZFP36L2*) and *Y* (*RHOB*). Pearson correlation = 0.465. **d**, The scatter plot depicting the correlation between residual of *X* (*ZFP36L2*) condition on *Y* (*RHOB*) and residual of *Y* (*RHOB*) condition on the ligand (*RPS19*) and receptor (*C5AR1*). Pearson correlation = 0.259. **e**, UMAP plots of *X* (*ZFP36L2*) and ligand (*RPS19*) in T cells, and UMAP plots of *Y* (*RHOB*) and receptor (*C5AR1*) in myeloid cells. **f**, An illustration of the CCC network between T cells and myeloid cells, represented using GEMs and ligand-receptor pairs. The size of each circle reflects the number of DEGs within each GEM. The composition of DEGs in GEM_T_10 and GEM_M_10 is highlighted in this figure.

For each *X* and *Y* pair that passed the IV test, we systematically searched for potential LR pairs that likely transduce signals between cells by identifying the LR pairs using the conditional independence test (CIT). As an example, *ZFP36L2* is statistically correlated with *RHOB* (Pearson correlation = 0.465). It is interesting to notice that *RPS19* (*L*) and *ZFP36L2* (*X*) are co-expressed in the exhausted T cells; on the other hand, the receptor C5AR1 (*R*) and RHOB (*Y*) are co-expressed in macrophages (Figure 4e). After conditioning on the estimated interaction state of *RPS19*-*C5AR1*, *ZFP36L2* (*X*) and *RHOB* (*Y*) are much less dependent, with a Pearson correlation less than 0.3 (Figure 4d). This suggests that *RPS19*-*C5AR1* may be involved in the signal transduction between the changed state of T (reflected by *ZFP36L2*) and the changed state of myeloid cells (reflected by *RHOB*).

When anti-PD-1 treatment induces a change in the state of a signaling pathway in T cells, it often leads to multiple DEGs (multiple *X*s); similarly, a change in the state of a pathway in a downstream non-T cell induced by CCC can lead to multiple DEGs (multiple *Y*s). Thus, if a pair of LR is detected to underlie the correlation of a module of *X*s and a module of *Y*s, it would more likely represent a signal transduction between pathways in two types of cells than correlations between a single triplet of *X-LR-Y*. We set out to search for the *X-LR-Y* triplets that passed the IV and CIT tests and share a common LR. We hypothesize that these gene expression modules (GEMs) in the PD1+ T and corresponding non-T cells represent the state of a signaling pathway in respective cell types, and the LR pair mediates the transduction of signal from T to non-T cells. Thus, such a module of *X-LR-Y* triplets represents a CCC channel.

We systematically searched for triplet modules (Supplementary Figure 4a). As an example, we discovered that ligand-receptor pairs of *RPS19* and *C5AR1* potentially mediate correlated expression of GEM in T (named as GEM_T_10) and a GEM in myeloid cells (named as GEM_M_10) (Figure 4f). GEM_T_10 and GEM_M_10 are co-expressed with *RPS19* and *C5AR1* on UMAP plots (Supplemental Figure 4b). Before CIT, GEM_T_10 is significantly correlated with GEM_M_10 (Pearson correlation = 0.464, p-value < 0.001). After CIT, the correlation reduced significantly (Pearson correlation = 0.238) (Supplemental Figure 4c).

To investigate the pathways that potentially regulate the GEM, we used cGSA (manuscript under review; no preprint available), a contextual gene set analysis (GSA) pipeline designed to integrate experimental contexts into the pathway analysis of DEGs by using large language models (LLMs). We found that the communication patterns are distinct between E tumors and NE tumors (Supplemental Figure 5a, b). For example, in NE tumors only, we identified multiple potential causal relationships (*X-LR-Y* triplets) connecting DEGs in PD1+ T cells pathways to the myeloid DEGs involved in fatty acid metabolic processes. The fatty acid metabolic process in myeloid cells is well known to promote tumorigenesis and contribute to treatment resistance^21,22^. Therefore, this communication between PD1+ T cells and myeloid cells in NE tumors may provide insights into potential cell-cell communications involved in the resistance mechanism of anti-PD1 treatment.

### CD4+ and CD8+ exhausted T cells have significantly different CCC networks

We noted that CD4+ and CD8+ exhausted T cells utilized different ligand-receptor interactions to transmit various signals to other types of cells (Supplementary Table S2 and S3). Tumor-associated macrophages (TAMs), composed of anti-tumor M1-like and pro-tumor M2-like subsets, are key regulators of the tumor immune microenvironment and significantly influence the efficacy of anti-PD1 therapy^23^. Our findings indicate that CD4+ exhausted T cells communicate with M1-like TAMs to enhance their anti-tumor function through ligand-receptor interactions, including TNF-TNFRSF14^24^ and TNFSF14-LTBR^25^ (Figure 5a). Integrating the result of cGSA pathway enrichment analysis and causal inference analysis, we observed that the activation of signaling pathways in CD4+ exhausted T cells is causally linked to the activation of TNF-alpha signaling via NF-kB pathway in M1 TAMs through ligand-receptors (Figure 5b). Similarly, our results suggest that CD8+ exhausted T cells might engage in crosstalk with M1-like TAMs, modulating pathway states through ligand-receptor pairs such as TNF-TNFRS1A^26^, ICAM1-ITGAL/ITGB2^27,28^, and CCL8-CCR2^29^ to create an anti-tumor TME (Figure 6a, b).

**Figure 5.**
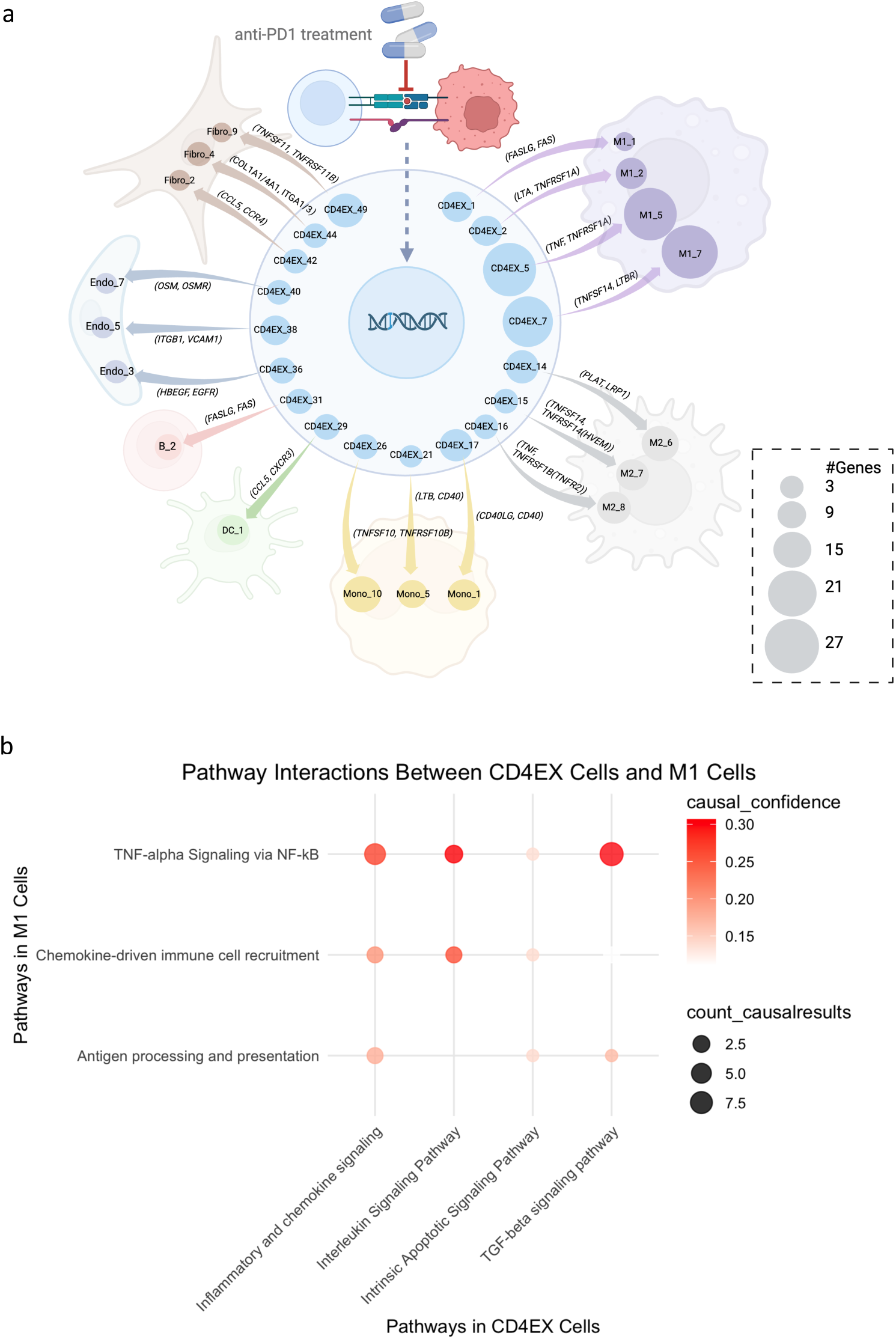
Summary of CCC network CD4+ exhausted cells with non-T cells in TME. **a**, The CCC network of CD4+ exhausted cells with non-T cells, including monocytes, B cells, dendritic cells, M1-like TAMs, M2-like TAMs, Fibroblasts, and endothelial cells. The size of each circle reflects the number of DEGs within each GEM. **b**, A dot plot of pathway interactions between CD4+ exhausted T cells and M1-like TAMs. The pathways are enriched using the cGSA method. The size of the dots indicates the count of causally related DEG pairs *X*-*Y* (*X* from T cells and *Y* from Myeloid cells). The color intensity of the dots indicates the significance of the causal relationship, which is the P-value of the Fisher-z CIT test.

**Figure 6.**
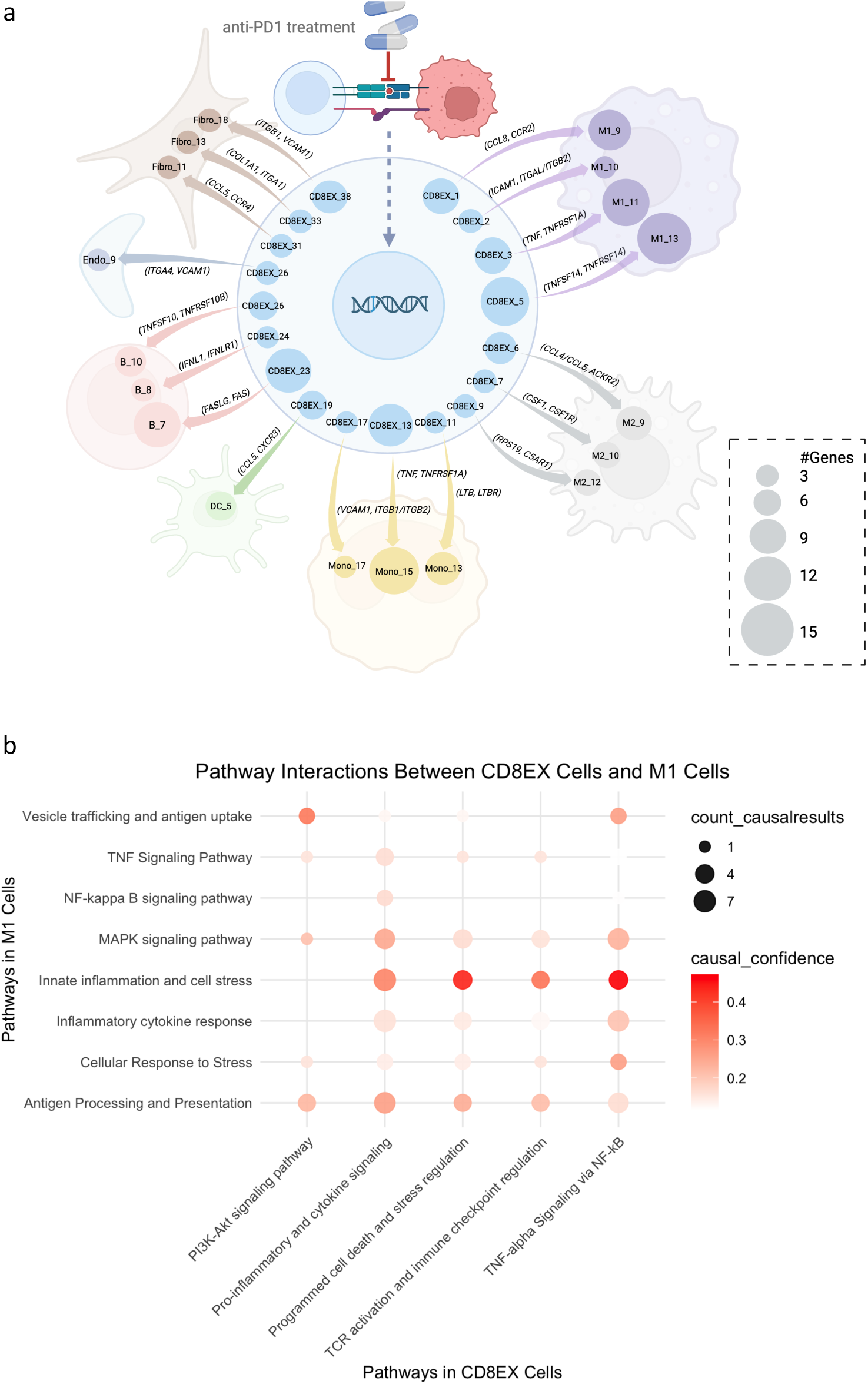
Summary of CCC network CD8+ exhausted cells with non-T cells in TME. **a**, The CCC network of CD8+ exhausted cells with non-T cells. **b**, A dot plot of pathway interactions between CD8+ exhausted T cells and M1-like TAMs.

Analysis of our dataset revealed that 89% of M1-like TAMs originated from E tumors, whereas 90% of M2-like TAMs were derived from NE tumors. CCCs between CD4+/CD8+ exhausted T cells and M1-like TAMs are mainly observed in the E tumors. Meanwhile, the CCCs between CD4+/CD8+ exhausted T cells and M2-like TAMs are mainly observed in NE tumors. Given that previous studies have established TCR clonotype expansion as a biomarker of responsiveness to anti-PD1 therapy^13^, our findings suggest that the CCC patterns from CD4+/CD8+ exhausted T cells to M1 TAMs are highly enriched in E tumors, providing mechanistic insights into anti-PD1 efficacy. Conversely, CCC between CD4+/CD8+ exhausted T cells and M2-like TAMs may elucidate how pro-tumor pathways are activated in NE tumors, contributing to resistance to anti-PD1 therapy. These CCC interactions represent potential therapeutic targets for enhancing responses to immune checkpoint blockade^30^.

Our data analysis suggests that after anti-PD1 treatment, CD4+/CD8+ may interact with M2 in the NE tumors. Among these ligand-receptors, some are targeted in clinical studies, so our study can “rediscover” the therapeutic strategies discovered by other approaches. For example, there are medications targeting TNFRSF1B (TNFR2)^31^, CSF1R^32,33^, and C5AR1^34,35^, and our results showed these receptors were likely involved in CCCs between T and M2 cells. Such results illustrate the potential utility of our framework for discovering CCC channels as future targets. Beyond established targets, several ligand-receptor interactions identified in our study hold potential for future immunotherapeutic development. TNFRSF14 (HVEM)^36^, CSF2RB^37^, and LRP1^38^ have been implicated in the regulation of M2-like TAMs and immune responses following anti-PD1 therapy. From the causal inference perspective, our CCC result shows that these receptors can be the next promising immunotherapy targets, combined with anti-PD1 treatment to improve the treatment efficiency.

## Discussion

Many studies have proved that various cell components of the TME communicate and influence each other ^39–41^. However, few studies have examined CCC among cells in a causal manner^42^. In our study, we analyzed publicly available data from breast cancer patients receiving anti-PD-1 treatment to reveal aspects of the CCC causal network in the TME. After anti-PD1 treatment, we observed DEGs not only in PD1+ T cells but also in non-T cells. Based on this result, we hypothesized that CCC involving PD1+ T cells drives these changes in other cells. We designed a process incorporating causal inference methods to evaluate this hypothesis and construct CCC networks in the TME triggered by anti-PD-1 therapy.

While it is possible to detect the correlation of certain genes across cell types, it is difficult to determine whether such a correlation is due to the CCC. Our approach used the treatment as an instrumental variable and employed IV analysis to eliminate possible latent confounding factors, providing evidence for CCC. Identification of ligand-receptor pairs that likely mediate the signal transduction in the channels discovered by IV analysis further provide mechanistic insights for potential CCC^43,44^. Finally, we construct a CCC network reflecting the overall impact of anti-PD1 treatment. Causal methodologies enable the identification of cell-type-specific CCC channels that would be difficult to achieve by other conventional single-cell analysis.

Our data analysis suggests that after anti-PD1 treatment, CD4+/CD8+ may interact with M2 in the NE tumors. Among these ligand-receptors, some are already targeted in clinical studies, so our study can “rediscover” the therapeutic strategies discovered by other approaches. For example, there are medications targeting TNFRSF1B (TNFR2)^31^, CSF1R^32,33^, and C5AR1^34,35^, and our results showed these receptors were likely involved in CCCs between T and M2 cells. Such results illustrate the potential utility of our framework for discovering CCC channels as future targets. Beyond established targets, several ligand-receptor interactions identified in our study hold potential for future immunotherapeutic development. From a causal inference perspective, our CCC results show that these receptors may be the next promising immunotherapy targets, combined with anti-PD1 treatment to improve the treatment efficiency. Furthermore, causal methods can be utilized to simulate the effect of perturbing a specific CCC channel. Thus, the causal discovery framework can serve as a powerful tool for computational drug discovery.

In summary, we demonstrated the utility of causal inference methods to investigate CCC in breast cancer patients receiving anti-PD-1 therapy. This analytical framework can be generalized to other single-cell RNA sequencing datasets to systematically elucidate causal CCC networks across various cancer types and treatments. In future studies, integrating causal inference with spatially resolved omics technologies, such as spatial transcriptomics, could substantially enhance precision by leveraging the spatial context of cellular interactions. Spatial omics would enable a deeper understanding of proximity-dependent communication, thereby refining causal network reconstruction and potentially revealing novel therapeutic targets. Moreover, extending these causal models to simulate pharmacological or genetic perturbations in specific CCC channels could accelerate computational drug discovery. Ultimately, this integrated causal-spatial approach holds significant promise for identifying robust biomarkers of immunotherapy response and uncovering innovative targets to improve clinical outcomes in cancer immunotherapy.

## Methods

### Data collection and primary process of single-cell data

We collected the single-cell sequencing data and TCR-sequencing data of 31 early-stage breast cancer patients from the study NCT03197389. This cohort includes 13 TNBC, 3 ER^-^/HER2^+^, and 15 ER^+^/HER2^+/-^ patients. Each patient has both before and after anti-PD-1 treatment. We used Scanpy^45^ to process single-cell data. After the aggregation and normalization, we removed the cells expressing fewer than 500 genes or more than 5,000 genes and the cells containing fewer than 400 UMIs (unique molecular identifiers) or more than 25,000 UMIs. Then, we filtered the cells with a high percentage (>20%) of mitochondrial genes. After filtration, we got 134,752 cells in total. We used unsupervised cell clustering methods for feature reduction. Cells were divided into 22 clusters and then annotated as five cell types according to the cell type markers expression levels.

### Identify differentially expressed genes

To identify differentially expressed genes (DEGs), we analyzed paired gene expression data from 31 patients, each with samples collected before and after treatment. Single-cell RNA sequencing data were aggregated into pseudo-bulk profiles and normalized using log2(TPM + 1) transformation. To account for inter-patient variability, we performed paired t-tests for each gene, comparing expression levels between the two conditions within each patient. Genes with a p-value < 0.05 and an absolute log2 fold-change greater than 1 were considered significant. To control for multiple testing, we applied the q-value method to adjust for the false discovery rate (FDR), identifying DEGs based on a stringent q-value threshold.

### Using instrumental variable (IV) analysis to remove the effect of confounders

We used the R package ivreg (version 0.6-3) for instrumental variable (IV) analysis to control for confounders. IV is implemented by using two-stage least-squares (2SLS) estimation. ^46^ In the first stage, we regress the endogenous variable *X* on the instrumental variable *Z* to obtain the predicted values ̂*X*.

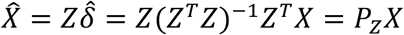

Where ̂𝛿 is the estimated coefficient from the regression of *X* on *Z.* PZ is the projection matrix formed from *Z*. After this step, ̂*X* is correlated with *Z* but not correlated with the error of this regression. In the second stage, we use the ̂*X* to perform the regression to *Y*.

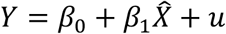

Where 𝛽_0_ and 𝛽_1_ are estimated coefficients, and 𝑢 is the error.

To apply IV analysis, three conditions must be met: relevance, exogeneity, and exclusion restriction. The instrumental variable *Z* must be correlated with the endogenous explanatory variable (X), uncorrelated with the outcome variable *Y* except through *X*, and uncorrelated with any confounders between *X* and *Y*. In our study, *Z* represents the anti-PD-1 treatment, *X* refers to the DEGs in PD1+ T cells or their subtypes, and *Y* corresponds to other cell-type DEGs. Since the treatment directly targets PD1+ T cells, *Z* is correlated with *X*. We calculated the residuals of *Y* regressed on *X* and filtered *X-Y* pairs where *Z* was uncorrelated with the residuals of *Y*, indicating that *Z* only affects *Y* through *X*. After confirming that our data met the IV assumptions, we grouped all cells by sample_id to generate pseudo-bulk data. We created a matrix with the expression levels of DEGs from both cell types (DEG *X* and *Y*) as columns and sample_ids as rows, adding a treatment column (0: before treatment, 1: after treatment). Applying ivreg to this matrix, we collected p-values for the IV test and filtered out DEG pairs with p-values greater than 0.05.

### CIT using the Fisher-z test

Fisher-z is a CIT method used to test the conditional independence between variables *X* and *Y*, given another variable *Z*^47^ (*X* ╨ *Y* | *Z*). In the fisher-z method, the correlation coefficient r between *X* and *Y* conditioned on *Z* is converted to a z-score using this equation:

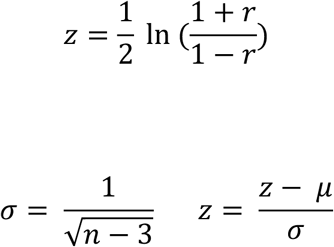

The null hypothesis is *X* and *Y* are independent, conditioning on *Z*. In the equation, 𝜇 is the mean of the fisher-z transformed distribution under the null hypothesis, and n is the sample size. Then, the p-value is calculated by comparing the *z-*score to the standard normal distribution.

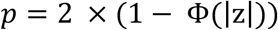

𝛷(|𝑧|) is the cumulative distribution function of the standard normal distribution, *|z|* is the absolute value of *z*. Because the purpose is to prove the null hypothesis is correct, we use p-value > 0.05 as the threshold to filter the results.

### Search for the ligand-receptor pairs used for cross-cell type communication

To investigate cross-cell type communication, we applied Fisher-Z conditional independence test (CIT) methods from the causal-learn Python package (https://github.com/py-why/causal-learn) to identify ligand-receptor pairs involved in cell-cell communication (CCC). We curated a dataset of 12,021 ligand-receptor pairs from KEGG^48^, Ramilowski^43^, PPI_prediction^49^, PPI_prediction_GO, and Guide2Pharmacology^50^ databases. As input data, we selected differentially expressed gene (DEG) pairs that passed the instrumental variable (IV) test (q-value < 0.05). CIT methods were then applied, conditioning on each ligand-receptor pair. In our model, we defined T cell DEGs’ pseudo-bulk expression levels as the causal variable *(X)*, other cell type DEGs’ pseudo-bulk expression levels as the effect variable *(Y)*, and pseudo-bulk expression levels of ligand multiplied by that of its receptor as the conditioning variable *(Z)*. During the search, we further constrained that ligands and receptors should be co-expressed with *X* and *Y* at the single-cell level, respectively (Pearson correlation > 0.3).

### Group the results to GEMs to build the CCC networks

We grouped the DEG pairs X and Y based on their involvement in the same ligand-receptor pairs to establish CCC. The results were filtered by applying a Pearson correlation threshold of less than 0.3 for the following comparisons: between *X* and *Y*, *X* and the ligand, *X* and the ligand-receptor complex, the receptor and, and the ligand-receptor complex (LR) and *Y*. The co-expressed and co-regulated DEGs were then grouped into gene expression modules (GEMs). To validate our findings, we conducted a literature review to find the results that were supported by existing studies. Finally, we used BioRender (https://app.biorender.com/) to organize the results and create visual representations of the cell-cell communication networks in response to anti-PD-1 treatment.

### Overview of cGSA workflow

Contextual gene set analysis (cGSA) integrates experimental context into pathway analysis of DEGs using large language models (LLMs). By incorporating user-provided background information, cGSA enhances pathway relevance and assigns a contextual relevance score to each pathway. The pipeline consists of four steps: (1) detecting gene clusters using the EdMot algorithm on DEG networks from STRING, (2) performing enrichment analysis via GSEA, (3) screening pathways with LLMs to refine relevance based on experimental context, and (4) summarizing pathways with LLMs to highlight key findings.

We applied cGSA to DEGs from the cell types in 31 breast cancer patients undergoing anti-PD-1 therapy, using cell-type-specific contexts. Pathway enrichment was performed with KEGG, WikiPathways, Reactome, MSigDB, and Panther, with human as the reference species.

## Acknowledgments

This research was supported in part by The National Library of Medicine; National Cancer Institute at the National Institutes of Health grants R01LM012011, R00LM013089 and R01CA254274. The Department specifically disclaims responsibility for any analysis, interpretations or conclusions.

## Data Availability Statement

All data analyzed in this study were obtained from a publicly available dataset published by Bassez et al. (Nature, 2021). The single-cell RNA-seq data from pre- and post-treatment breast cancer samples are available through the European Genome-phenome Archive (EGA) under accession number EGAS00001004809.

## Notes

### Competing Interest Statement

The authors have declared no competing interest.

